# Is level-1 blob reconstruction under the network multispecies coalescent easy?

**DOI:** 10.64898/2026.06.06.730607

**Authors:** Junyan Dai, Erin K. Molloy

## Abstract

Hybridization is an important evolutionary process, commonly modeled by the network multispecies coalescent. Reconstructing evolutionary histories under this model is notoriously costly, even for level-1 networks where hybridization events are isolated from each other. The widely used methods that combine speed with statistical guarantees rely on quartet concordance factors computed for all subsets of four species, resulting in an *O*(*n*^4^*k*) bottleneck that severely limits scalability to large numbers of species (*n*) and genes (*k*). Among quartet-based methods, NANUQ+ is notable because it decomposes the problem into two steps: first reconstructing a *tree of blobs*, which compresses each non-treelike part of the network, called a blob, into a single vertex, and second reconstructing the internal structure of each level-1 blob, specifically its circular order and hybrid vertex. Here, we investigate whether level-1 blob reconstruction is difficult once the tree of blobs is known. We present a fast and statistically consistent algorithm, called NetCS, based on two simple primitives: majority voting and merge sort, circumventing the bottleneck of computing all quartet concordance factors. In simulations, NetCS achieved comparable accuracy to NANUQ+ and was dramatically faster, enabling analyses of 200 taxa and 1000 genes in only a few minutes. Both methods attained near-perfect accuracy when given the true tree of blobs; however, their performance degraded in end-to-end pipelines due to errors in tree of blobs reconstruction. Strikingly, even methods that reconstruct level-1 networks directly struggled to accurately predict hybrid ancestry. Our results suggest that reconstructing level-1 blobs is unexpectedly easy once the tree of blobs is known, and that a major challenge for phylogenetic network inference lies in accurate tree of blobs reconstruction. NetCS is implemented within TREE-QMC, available at https://github.com/molloy-lab/TREE-QMC

## 1 Introduction

Hybridization and admixture are important evolutionary scenarios that must be represented as phylogenetic networks rather than trees. A phylogenetic network governs the ancestry of unlinked, recombination-free genomic regions, called genes, commonly modeled by the Network Multispecies Coalescent (NMSC) [40, 41]. Reticulations in the network enable genes from the same species’ genome to trace back to different ancestral populations, resulting in conflicting genealogies, called gene trees. Additionally, conflicts among gene trees can arise from incomplete lineage sorting (ILS), similar to the traditional MSC model [29]. Although gene trees are not directly observed, they govern the evolution of DNA sequences, from which they can be estimated with fast methods, such as RAxML [36] and IQ-TREE [26].

Reconstructing phylogenetic networks under the NMSC is notoriously costly even for level-1 networks where hybridization events are isolated from each other. The popular methods that combine speed with statistical guarantees rely on *quartet concordance factors (qCFs)* computed for all subsets of four species, called taxa. A quartet is an unrooted phylogenetic tree on four species, and its concordance factor is simply the number of gene trees that have the same topology after restriction to the same species. Performing this precomputation for all four-taxon subsets results in an *O*(*n*^4^*k*) cost that severely limits scalability to large numbers of species (*n*) and genes (*k*), although methods based on qCFs are still significantly cheaper than those that based on full likelihood calculations (e.g., [41]).

The first network reconstruction method to leverage qCFs was SNaQ [34, 35], which conducts a search over the space of (semi-directed) level-1 networks via edit moves, seeking a network to maximize its pseudo-likelihood function (see [24] for a related composite likelihood function for DNA sequences). To improve scalability beyond ∼30 species, the most recent version of SNaQ attempts to approximate pseudo-likelihood by randomly sampling four-taxon subsets [22]; this approach currently has no statistical guarantees, and there are intricacies to random sampling, as highlighted by [10]. Alternatively, SNaQ can be leveraged within the divide-and-conquer framework InPhyNet [23], but then the goal is to run SNaQ on pairwise disjoint subsets of taxa that are as large as possible.

Another recently developed method that leverages qCFs is CAMUS [38], accepted to ISMB 2026. Unlike SNaQ, CAMUS takes a rooted base tree as input (typically estimated via ASTRAL-IV [43]) and then adds reticulations edges to maximize the number of quartets also displayed by the (directed) level-1 network (also see [28]). Although CAMUS is more scalable than SNaQ, it still requires qCF precomputation. CAMUS does not currently have statistical guarantees under the NMSC and can be statistically inconsistent when ASTRAL-IV is used to construct the base tree, as recently shown by Dinh and Baños [11] (similar issues for tree-based network inference arise in the context of admixture graph reconstruction [27]).

In contrast, NANUQ+ [4] decomposes (semi-directed) level-1 network reconstruction into two steps: (1) reconstructing a tree of blobs, which compresses each non-treelike part of the network, called a blob, into a single vertex and then (2) reconstructing the internal structure of each level-1 blob, specifically its circular order and hybrid vertex (also see [16]). Each step as implemented by NANUQ+ is provably *statistically consistent* under the NMSC with some assumptions [3, 4]; however, both steps require qCF precomputation, limiting scalability.

Recent work has focused on scalable tree of blobs reconstruction from qCFs with statistical and/or correctness guarantees (e.g., [3, 10, 17, 13, 30]). This motivates us to investigate whether level-1 blob reconstruction is difficult once the tree of blobs is known. The remainder of the paper is organized as follows. We present the terminology and background in Section 2. In Section 3, we present our new algorithm for level-1 blob reconstruction from qCFs, called NetCS, along with its theoretical properties. NetCS is based on two simple primitives: majority voting and sorting; thus it differs dramatically from NANUQ+’s approach, which computes distances between all pairs of taxa based on the input qCFs and then evaluates the goodness of fit between expected and observed distances for all possible level-1 blob structures. In Section 4, we describe the results of benchmarking NetCS against NANUQ+ and CAMUS. We conclude in Section 5 with a discussion of whether level-1 blob reconstruction is difficult based on our theoretical and empirical results.

## 2 Background

To present Net-CS, we review terminology and background for phylogenetic networks.

### Phylogenetic Networks

A *directed* phylogenetic network *N* ^+^ is a directed simple acyclic finite graph with the property that there exists a directed path from the root, a unique vertex with in-degree 0, to all other vertices. The leaf vertices of *N* ^+^, which have out-degree 0, are bijectively labeled by a set *S* of species. A *hybrid* is a vertex with in-degree *>* 1, and a *reticulation edge* is an arc whose child is a hybrid. A *wedge* path in *N* ^+^ is a sequence of distinct vertices *u*_0_, *u*_1_, …, *u*_*k*−1_, *u*_*k*_ such that there exists some vertex *u*_*i*_ that “roots” the path so *u*_*i*_, *u*_*i*−1_, …, *u*_2_, *u*_1_ and *u*_*i*_, *u*_*i*+1_, …, *u*_*k*−1_, *u*_*k*_ are both directed paths in *N* ^+^ [17]. A directed network *N* ^+^ can be converted into a *semi-directed* network *N* ^−^ [34] by removing all edges and vertices above the lowest stable ancestor *s* of *S* (i.e., the lowest vertex on all directed paths from the root to each leaf in *S*; see [37], page 263), undirecting all non-reticulation edges, and suppressing *s* if it has degree 2. The wedge path terminology is used for semi-directed networks because there is a bijection between wedge paths in *N* ^+^ and paths in *N* ^−^ that do not traverse two reticulation edges incident to the same hybrid vertex. If there are no hybrid vertices, *N* ^−^ is simply an unrooted phylogenetic tree. We assume that semi-directed networks and unrooted trees are *binary*, specifically leaves have degree 1 and all other vertices have degree 3.

### Cut Edges, Blobs, and Partitions

A bridge, also called a *cut edge*, is an edge whose deletion increases the number of connected components of the graph. The maximal bridgeless subgraphs of *N* ^−^, when ignoring the directionality of reticulation edges, are called *blobs*. The *level* of a blob is the number of reticulation edges in it minus the number of hybrid vertices in it; we assume that all blobs are level-1. The degree of a blob is the number of cut edges adjacent to it. An *m*-blob is a blob with degree *m*. The *tree of blobs (TOB)* [15] is obtained from *N* ^−^ by contracting the edges in each blob into a single vertex and then suppressing any degree-2 vertices (Fig. 1A–B). Let *v*_ℬ_ denote the vertex in the tree of blobs *T*_*N*_ that corresponds to blob ℬin the underlying network *N* ^−^. The deletion of vertex *v*_ℬ_ from *T*_*N*_ produces a collection of rooted subgraphs whose leaf sets partition the taxon set *S*, denoted 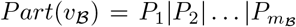, where *m*_ℬ_ is the degree of *v*_ℬ_; this is equivalent to the partition induced by deleting all edges and vertices in blob ℬfrom *N* ^−^, denoted *Part*(ℬ). Likewise, the deletion of a cut edge *e* from *N* ^−^ or its TOB yields two rooted subgraphs whose leaves partition *S* into two subsets, called a *bipartition* or *split*, denoted *A*|*B*. A bipartition is trivial if either |*A*| = 1 or |*B*| = 1. If both endpoints of *e* have degree 3, then deleting *e*, along with its endpoints, induces a *quadrapartition*: *Quad*(*e*) = *A*_1_|*A*_2_|*B*_1_|*B*_2_.

**Fig. 1:**
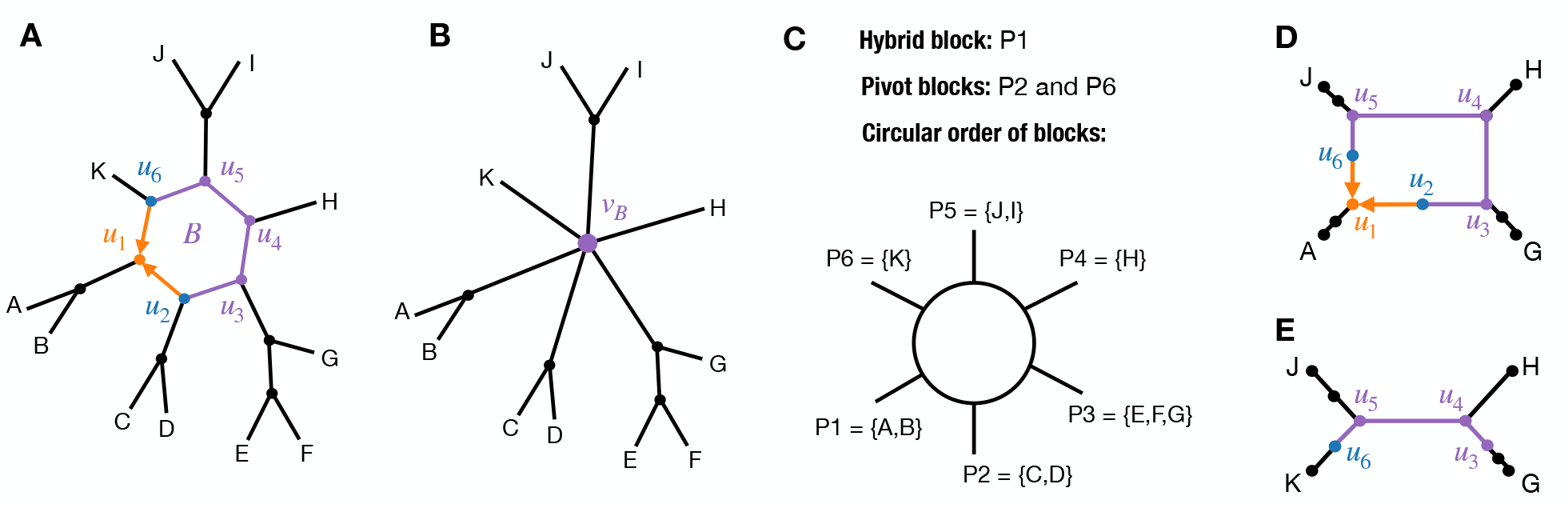
**A)** A binary, semi-directed, level-1 network *N* ^−^ on species set *S* with one non-trivial blob ℬ. **B)** Tree of blobs for *N* ^−^. Blob vertex *v*_ℬ_ induces an 6-partition of *S* denoted *Part*(*v*_ℬ_) = *P*_1_| … |*P*_6_. **C)** Information about ℬ. **D)** Subnetwork on 4 taxa from distinct blocks of *Part*(*v*_ℬ_), including hybrid block *P*_1_. The sub-network displays two quartets: *A, J*|*H, G* and *A, G*|*H, J*. **E)** Subnetwork on 4 taxa from distinct blocks of *Part*(*v*_ℬ_), excluding hybrid block *P*_1_.

### Blob structure

Each blob in a binary, semi-directed, level-1 network is either a single vertex or a cycle when ignoring the directionality of reticulation edges; the former is called trivial and the latter non-trivial. Moreover, no two blobs can share an edge or vertex, and each non-trivial blob contains exactly one hybrid vertex incident to exactly two reticulation edges. The **circular order** of ℬ is the cyclic ordering of vertices encountered when traversing the cycle ignoring edge direction, considered up to rotation and reversal. There is a bijection between the vertices in the circular order of blob ℬ and the blocks of *Part*(ℬ); thus, we refer to them interchangeably. *N* ^−^ can be reconstructed from its TOB by (1) determining the circular order of each non-trivial blob, and (2) identifying the hybrid of each non-trivial blob, as this gives the directionality of reticulation edges (Fig. 1C) [4].

### Quarnets and quartets

A *subnetwork* of *N* ^−^ on *Y* ⊆ *S*, denoted *N* ^−^|_*Y*_, is the subgraph induced by the union of vertices and edges on all wedge paths in *N* ^−^ between all pairs of taxa in *Y* [5]. The *restriction* of *N* ^−^ to *Y* ⊆ *S* is formed by computing the subnetwork on *Y* and then repeatedly suppressing degree-2 vertices, parallel arcs, and 2-blobs [17]. For |*Y*| = 4, subnetworks are referred to as **quarnets. Quarnet-splits** are the set of bipartitions induced by the cut edges of *N* ^−^|_*Y*_ or equivalently the edges in the TOB of *N* ^−^|_*Y*_ [17]. A quarnet either contains a 4-blob or not [3, 17]. In the former case, the TOB of *N* ^−^|_*Y*_ is an (unresolved) star tree (Fig. 1D), and in the latter case, the TOB of *N* ^−^|_*Y*_ is a (binary) **quartet** tree (Fig. 1E); thus, we say that the evolutionary relationship is “treelike” [3]. A quartet on species set {*a, b, c, d}* has one of three possible topologies, indicated by the one non-trivial bipartition: *a, b*|*c, d* or *a, c*|*b, d*, or *a, d*|*b, c*. Any quarnet that contains a 4-blob **displays** exactly two quartets formed by deleting each of the reticulation edges incident to the hybrid vertex given our assumption that *N* ^−^ is binary and level-1 [34, 7].

### Model and Data

The Network Multispecies Coalescent (NMSC) model is parameterized by *N*^+^ = (*N* ^+^, *S, Θ*), where *N* ^+^ is a directed phylogenetic network on species set *S* and *Θ* is a set of numerical parameters: branch lengths in coalescent units, reticulation edge inheritance proportions, and the inheritance correlation *ρ* (note that we assume independent inheritance, i.e., *ρ* = 0). We only briefly summarize the NMSC here and refer readers to [34,10] for a complete description. A gene genealogy is generated under the NMSC by sampling one gene per species and tracing ancestry backward in time until all lineages coalesce (i.e., trace back to a common ancestor), forming a rooted, binary tree on *S* [40, 12]. We let ℙ_*N*_ + (*t*) denote the probability of an unrooted, binary gene tree *t* on species set *Y* ⊆ *S* given *N*^+^. For 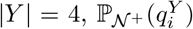, where *i* = {1, 2, 3} indexes the three possible quartets on *Y*, are the *expected* **quartet concordance factors (qCFs)**. Expected qCFs are fully determined by the semi-directed version of *N*^+^, denoted *N*^−^ (Proposition 3 in [34]). *Observed* qCFs are computed by counting the number of *k gene trees* whose restriction to *Y* is isomorphic to 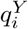, denoted 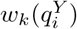, and then dividing each element in the 3-vector by its sum. Observed qCFs converge simultaneously to their expected values with probability 1 as the number of gene trees goes to infinity by the strong law of large numbers.

### Assumptions and Identifiability

We make standard assumptions about the model network; see [34, 7, 3, 4, 10]. Specifically, we assume that *N*^−^ is binary, semi-directed, level-1, and metric (i.e., non-reticulation edges have positive lengths, reticulation edges have lengths greater than or equal to zero, reticulation edges incident to the same target have inheritance proportions that sum to 1 and take on values between 0 and 1, non-inclusive). Under these assumptions, the structure of 3-blobs is <u>not</u> identifiable from qCFs [7]. However, the circular order of 4-blobs is identifiable, and for large blobs with degree *>* 4, the hybrid vertex also is identifiable (Theorem 4 in [7]). The crux behind this identifiability result is that for any 4-taxon subset *Y* such that *N* ^−^|_*Y*_ contains a 4-blob, the qCFs on *Y* are distinct, i.e., 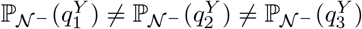 (Def. 17 and Theorem 3 in [7]). Moreover, the lowest CF corresponds to the only quartet that is <u>not</u> displayed by *N* ^−^|_*Y*_ (Proposition 11 in [7]). These two properties hold under generic numerical parameter settings except those in a set of measure zero (an example in the set of measure zero can be obtained by setting all edges in Fig. 1A to have the same length in coalescent units and both inheritance proportions to 0.5).

### Oracle for 4-blobs

Lastly, our approach requires an oracle that takes a 4-taxon subset on *Y* as input, specifically its associated 3-vector of qCFs, and returns whether the quarnet *N*^−^|_*Y*_ contains a 4-blob or not. In practice, we employ the same hypothesis testing framework used for tree of blobs reconstruction in NANUQ+ [4, 3, 2]. There are two versions of the hypothesis test: T3 and cut. The null hypothesis of both is that the *N* ^−^|_*Y*_ does <u>not</u> contain a 4-blob, thus low *p*-values indicate the null should be rejected (i.e., *N* ^−^|_*Y*_ does contain a 4-blob). The test assumes that expected qCFs are distinct for every 4-taxon subset *Y* ⊆ *S* such that *N* ^−^|_*Y*_ contains a 4-blob [3], which is satisfied as noted in the previous section. However, the T3 version of the hypothesis test additionally assumes that networks are *class-1 quartet-nonanomalous*, that is, for every 4-taxon subset *Y* ⊆ *S* such that *N* ^−^|_*Y*_ does <u>not</u> contain a 4-blob, 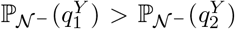 and 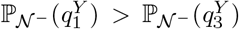, where 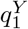 corresponds to the TOB of *N* ^−^|_*Y*_ [3]. This assumption is satisfied for common inheritance (*ρ* = 1) but not necessarily independent inheritance (*ρ* = 0) [5], although strong empirical performance of the T3 option is still possible even under model violations [3, 10].

## 3 Methods

We are now ready to present our method NetCS, which has the following input and output.

– **Input:**
  - Unrooted tree of blobs *T*_*N*_ on a set *S* of *n* species
  - Set {*G*_1_, …, *G*_*k*_ *}* of unrooted, binary gene trees, each on species set *S*_*i*_, where 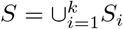
– **Output:** A binary, semi-directed, level-1 network on *S* with TOB isomorphic to *T*_*N*_ (note that only non-trivial blobs of degree at least 5 are reconstructed)

NetCS processes each vertex with degree 5 or more in *T*_*N*_ *independently* with two primitives: majority voting to detect the hybrid and sorting to resolve the circular order.

### 3.1 Hybrid Majority Voting

Consider some vertex *v*_ℬ_ in the input TOB *T*_*N*_ with degree *m*_ℬ_ ≥ 5. The first step in NetCS is to find the hybrid block of *Part*(*v*_ℬ_), which is the unique block induced by the cut edge incident to the reticulation vertex. Our approach is based on the following result.

#### Lemma 1.

*Let N*^−^ = (*N* ^−^, *S, Θ*) *be a binary, semi-directed, level-1, metric species network. Let O*_*k*_(*Y* ) *denote the p-value of NANUQ+ hypothesis test applied to some 4-taxon subset Y* ⊆ *S, specifically the qCFs computed from k independent (unrooted) gene trees on species set S sampled from the distribution induced by the NMSC given N* ^−^. *Let v*_ℬ_ *be a vertex in the TOB for N* ^−^ *with degree m*_ℬ_ ≥ 5. *Now consider two 4-taxon subsets constructed such that every taxon is drawn from distinct blocks of Part*(*v*_ℬ_), *with the first subset Y*_+*h*_ *drawing one taxon from the hybrid block of Part*(*v*_ℬ_) *and the second subset Y*_−*h*_ *drawing zero taxa from the hybrid block. Then, with probability* → 1 *as k* → ∞, *O*_*k*_(*Y*_+*h*_) *< O*_*k*_(*Y*_−*h*_) *for all pairs of 4-taxon subsets Y*_+*h*_, *Y*_−*h*_ *constructed as described above*.

*Proof*. For every pair of 4-taxon subsets *Y*_+*h*_, *Y*_−*h*_ constructed as described above, 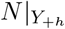 contains a 4-blob but 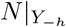 does not, by Lemma 5 in [10]. The remainder of the proof follows from the statistical consistency of the hypothesis test *O*_*k*_; specifically, Theorem 4 in [3] states that there exists a sequence of significance thresholds *α*_*k*_ →0 such that the hypothesis test applied to *k* gene trees will, with probability tending to 1 as *k* → ∞, return the correct result for all 4-taxon subsets of *S* (also see [30]). This implies the returned *p*-value will be less than *α*_*k*_ for any 4-taxon subset *Y* such that *N*|_*Y*_ contains a 4-blob and will be greater than or equal to *α*_*k*_ otherwise, giving us our result.

The innovation behind NetCS is applying the hypothesis test *O*_*k*_ to carefully selected 4-taxon subsets so that the precomputation of qCFs for all 4-taxon subsets of *S* can be avoided. Specifically, we create *re-quadrapartitions* of *Part*(*v*_ℬ_) and then apply *O*_*k*_ to four taxa drawn from distinct blocks.

#### Definition 1

(Re-quadraparition). *Let P* = *P*_1_|*P*_2_|…|*P*_*m*_ *be a m-partition with m* ≥ 5. *A* re-quadrapartition *is a coarsening of P into exactly 4 blocks, denoted* 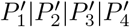, *such that each original block P*_*i*_ *is contained entirely within exactly one new block* 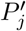.

Given a re-quadrapatition of *Part*(*v*_ℬ_), denoted 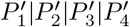, the hypothesis test *O*_*k*_ is applied within the **3-fix, 1-alter (3f1a) algorithm**, introduced at RECOMB 2026 [10]:

1. Select one taxon arbitrarily from each block of the requadrapartition: 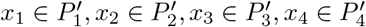.
2. Apply *O*_*k*_ to the qCFs for {*x*_*a*_, *x*_2_, *x*_3_, *x*_4_} for all 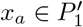.
3. Repeat step (2) three times but instead of fixing the three taxa from 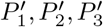 and rotating through all taxa in 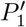, swap out 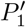 with 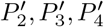.
4. Return the minimum *p*-value found from all applications of *O*_*k*_, along with the associated minimizer 4-taxon subset.

Unlike its original application, we leverage 3f1a for voting, allowing each taxon in the minimizer subset vote for its respective block of *Part*(*v*_ℬ_). This process is repeated for *c* rounds, each corresponding to an arbitrary re-quadrapartition of *Part*(*v*_ℬ_). The block of *Part*(*v*_ℬ_) with the most votes is selected as the hybrid, with some additional steps for tie breaking (Algorithm 1); the tie breaking step is closely related to hybrid identifiability [7, 13].

#### Theorem 1.

*Assume the same settings as Lemma 1, and let v*_ℬ_ *be some vertex in the TOB for N* ^−^ *with degree m*_ℬ_ ≥ 5. *Then, Algorithm 1 with c* ≥ 1 *voting rounds applied to Part*(*v*_ℬ_) *and k gene trees will, with probability* → 1 *as k* → ∞, *return the (correct) hybrid block of Part*(*v*_ℬ_). *The time complexity of applying Algorithm 1 to all v*_ℬ_ *is O*(*n* log (*n*)*k*) *in fast mode or O*(*n*^2^*k*) *otherwise, where n is the number of taxa*.

*Proof*. Vertex *v*_ℬ_ is associated with a non-trivial blob ℬ in *N* ^−^ given our assumption that *N* ^−^ is level-1 and binary, so the 3f1a algorithm applied to an arbitrary re-quadrapartition of *Part*(*v*_ℬ_) is guaranteed to apply *O*_*k*_ to at least one 4-taxon subset *Y* such that *N* ^−^|_*Y*_ contains a 4-blob, implying that *Y* contains a taxon from the hybrid block, by Theorem 2 in [10]. Then, by Lemma 1, the minimizer subset returned by 3f1a given an arbitrary re-quadrapartition of *Part*(*v*_ℬ_) and *k* gene trees will, with probability going to 1 as *k* → ∞, include the hybrid block of *Part*(*v*_ℬ_). Repeating for *c* arbitrary re-quadrapartitions of *Part*(*v*_ℬ_) will produce one of two outcomes: (1) the hybrid block is the only block that earns *c* votes or (2) the hybrid block is tied for *c* votes with one or more other blocks. Taking the five blocks with the most votes, breaking ties arbitrarily, ensures that exactly one way of selecting 4 out of these 5 blocks does not include the hybrid block. The *p*-value from applying the oracle *O*_*k*_ to this special 4-block subset will be <u>higher</u> than the other 4-block subsets by Lemma 1, revealing the hybrid block by exclusion. The argument above extends to *fast mode*, in which a single taxon is selected arbitrarily to represent each block of *Part*(*v*_ℬ_) (Algorithm 1, line 3), as any 4-taxon subset that covers the same distinct blocks of *Part*(*v*_ℬ_) yields the same expected qCFs under the NMSC given *N*^−^ [7] (also see [10]). It is worth noting that a 4-block tie occurs in fast mode because of setting *c* = 1. However, our implementation of NetCS uses *c* = *m*_ℬ_ −3, as this does not increase the total time complexity of NetCS (Corollary 1).

Now we consider time complexity. Building the taxon-to-block mapping function *F* is *O*(*m*_ℬ_) in fast mode or *O*(*n*) otherwise. Applying 3f1a to *c* re-quadrapartitions requires *O*(*cm*_ℬ_) or *O*(*cn*) queries to the oracle *O*_*k*_, depending on whether fast mode is employed or not. The oracle is a hypothesis test that can be performed in

#### Algorithm 1

Hybrid Majority Voting Algorithm

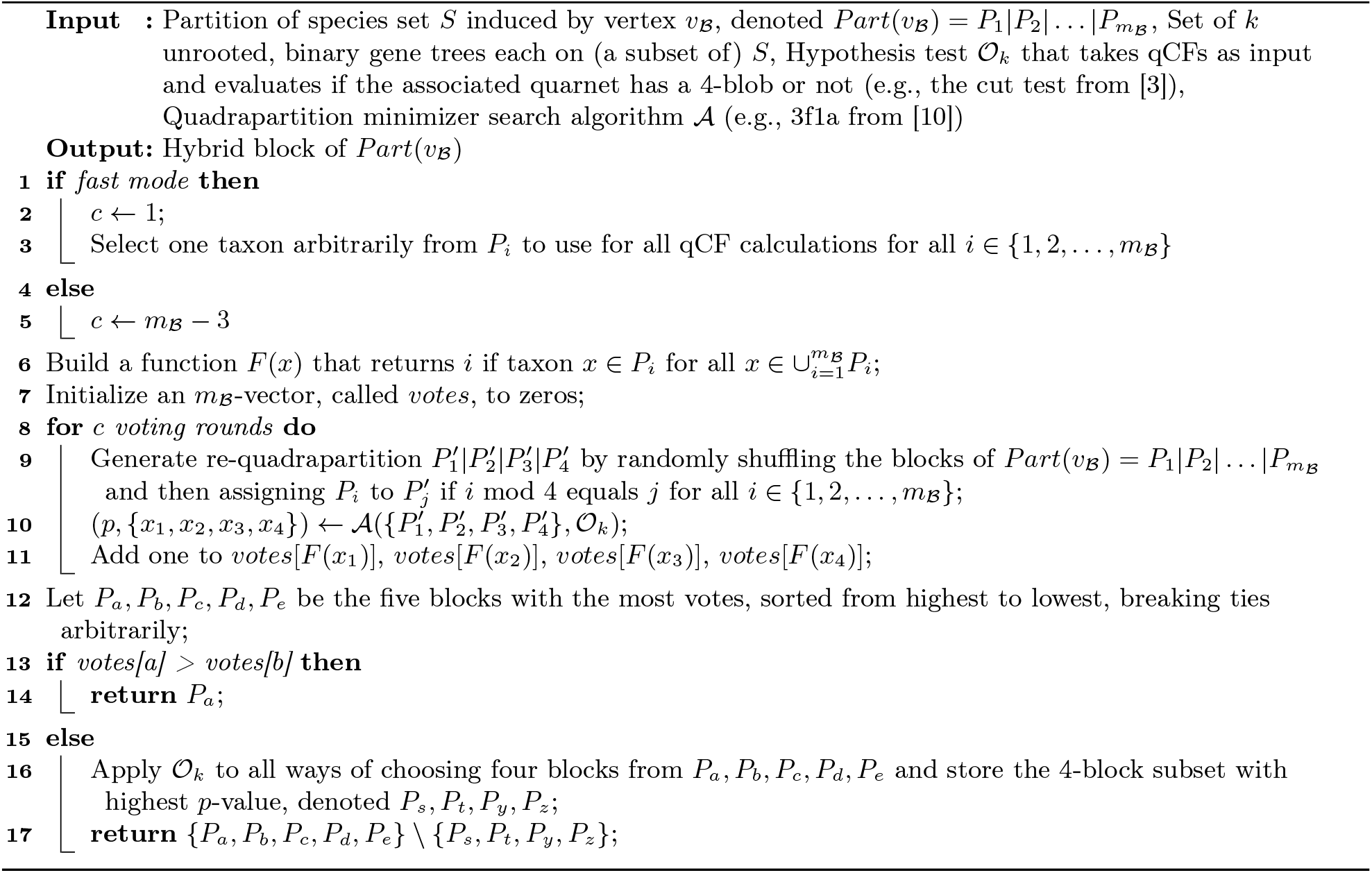

constant time but requires qCFs as input. After an *O*(*n* log (*n*)*k*) preprocessing step, the well-known binary lifting algorithm can be used to retrieve the lowest common ancestor of any pair of leaves in an (arbitrarily) rooted binary gene tree in constant time [14], enabling qCF computation for any 4-taxon subset on the fly in *O*(*k*) time. Therefore, when including the calculation of qCFs, each oracle query is *O*(*cm*_ℬ_*k*) in fast mode and *O*(*cnk*) otherwise. The other calculations do not exceed these time costs. Repeating for all *v*_ℬ_ gives 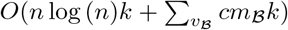 in fast mode and 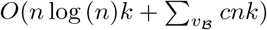 otherwise. Taking *c* = 1 in fast mode and *c* = *O*(*m*_ℬ_) otherwise gives us our result, noting that 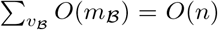. Note that hybrid detection requires qCFs to be computed for *O*(*n*) 4-taxon subsets when running in fast mode; as expected qCFs reveal the quarnet topology, we say it makes *O*(*n*) **quarnet queries**.

### 3.2 Pivot Scanning

After determining the hybrid block of *Part*(*v* _ℬ_), it remains to determine the circle order. NetCS leverages a *pivot* to linearize the circular order for sorting.

#### Definition 2

(Pivots). *Let N* ^−^ *be a binary, semi-directed, level-1 network. Then, any non-trivial blob ℬ in N* ^−^ *has two reticulation edges whose source vertices are called* pivots.

The deletion of the cut edge incident to each pivot vertex induces a pivot block.

#### Lemma 2.

*Let* 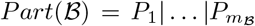 *be a partition induced by a non-trivial blob* ℬ*in a binary, semi-directed, level-1 network N* ^−^, *and let P*_*h*_ *be the hybrid block of Part*(ℬ). *Then, P*_*p*_ *is a pivot block of Part*(ℬ) *if and only if for all s, t* ∈ {1, …, *m* _ℬ_} *such that s*≠ *t* ≠ *p* ≠ *h and all* 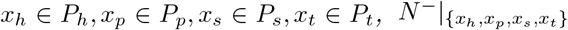 *displays quartet x*_*h*_, *x*_*p*_|*x*_*s*_, *x*_*t*_.

The proof of Lemma 2 is in the Appendix A. The idea is that the taxa from the pivot and hybrid blocks must be siblings in one of the two quartets displayed by the quarnet. As the lowest of the three qCFs corresponds to the non-displayed quartet by Proposition 11 in [7], the qCFs identify one block, out of four, that is neither the hybrid nor one of the two pivots. Then, scanning through all blocks in *Part*(*v* _ℬ_) gives the two pivot blocks by process of elimination (Algorithm 2 in Appendix A). This approach requires a method ℳ_*k*_ for computing the qCFs based on four distinct blocks of *Part*(*v* _ℬ_). In this setting, taxa from the same block are exchangeable, as previously noted, so in fast mode, one taxon is arbitrarily selected from each block in {*P*_*h*_, *P*_*p*_, *P*_*s*_, *P*_*t*_*}* for qCF computation; however, computing the average qCFs from all 4-taxon subsets covering {*P*_*h*_, *P*_*p*_, *P*_*s*_, *P*_*t*_*}* may increase robustness to noise, which occurs when qCFs are computed from a finite number of gene trees with estimation error and/or missing taxa. Both approaches enable consistent estimation.

#### Theorem 2.

*Assume the same settings as Lemma 1. Let π be the error tolerance:*

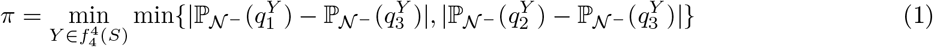

*where* 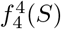 *is the set of all 4-taxon subsets Y* ⊆ *S such that N* ^−^|_*Y*_ *contains a 4-blob and* 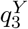 *is the quartet out of the three possible on Y that is <u>not</u> displayed by N* ^−^|_*Y*_ . *Let v* _ℬ_ *be some vertex in the TOB for N* ^−^ *with degree m* _ℬ_ ≥ 5. *Then, for every ϵ >* 0, *Algorithm 2 applied to Part*(*v* _ℬ_), *with hybrid block P*_*h*_, *and* 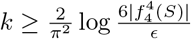 *gene trees returns the (correct) two pivot blocks of Part*(*v*_ℬ_ ) *with probability at least* 1 − *ϵ. The time complexity of applying Algorithm 2 to all v* _ℬ_ *is O*(*nk*) *in fast mode and O*(*n*^2^*k*) *otherwise*.

*Proof*. By Lemma 2, Algorithm 2 returns the two pivot blocks for all *v*_ℬ_ when the lowest qCF on *Y* corresponds to the non-displayed quartet for all 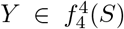 such that *N* ^−^|_*Y*_ contains a 4-blob. In fast mode or otherwise, this occurs when the error in the observed qCFs is bounded by *π/*2, specifically: 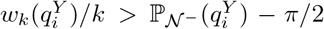 and 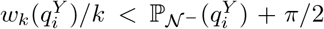 for all *i* = {1, 2, 3} and all 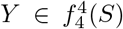. Following the approach in [33], let 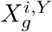 be a random variable which takes on value 1 when the restriction of gene tree *G*_*g*_ to *Y* is isomorphic to 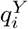 and takes on value 0 otherwise. Then, 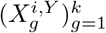 is a sequence of independent Bernoulli random variables with success probability 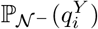, and the observed qCF is 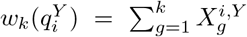 with expected value 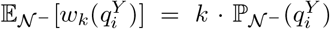. Now, we can bound the probability that 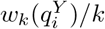 deviates from its expected value by *π/*2 or more using Hoeffding’s in-equality: 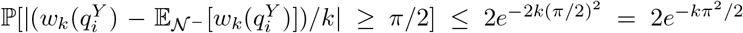 for all *i* ∈ {1, 2, 3} and all 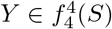. Let *E*_*i,Y*_ denote the event that 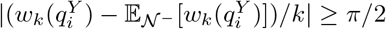. Then, by the union bound, 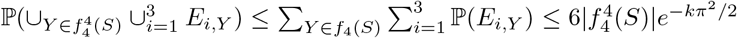. Lastly, we solve 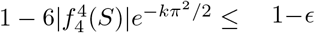 for *k* to get our result. To summarize, Algorithm 2 is consistent under the NMSC and, as 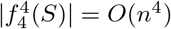, has sample complexity 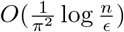.

Now consider time complexity. Algorithm 2 computes qCFs for *O*(*m*_ℬ_) 4-block subsets. Each qCF computation takes *O*(*k*) time in fast mode, taking advantage of the preprocessing performed for hybrid detection; otherwise, computing averages via the algorithm in Fig. S14 of [32] takes *O*(*nk*) time. Repeating for all *v*_ℬ_ gives 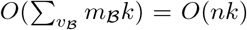 and 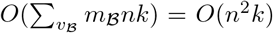. Pivot scan makes *O*(*n*) quarnet queries in fast mode.

### 3.3 Circular Order Sorting

Taking the hybrid block *P*_*h*_ and a pivot block *P*_*p*_ together enables us to linearize the circular order of blob ℬ as the canonical vector *V*_ℬ_ (*P*_*h*_, *P*_*p*_) = [*P*_*h*_, *P*_*p*_, …, *P*_*p*_*′* ], where *P*_*p*_*′* is the block of *Part*(ℬ) induced by the other pivot. Then, mapping each block *P*_*s*_ ∈ *Part*(ℬ) to its index in the canonical vector *V*_ℬ_ (*P*_*h*_, *P*_*p*_) yields a total ordering on the blocks, denoted *L*_ℬ_ (*P*_*h*_, *P*_*p*_). As an example, traversing the only cycle in *N* ^−^ of Fig. 1A from the hybrid vertex *u*_1_ towards the pivot *u*_2_, ignoring the direction of reticulation edges, gives *V*_ℬ_ (*P*_1_, *P*_2_) = [*P*_1_, *P*_2_, *P*_3_, *P*_4_, *P*_5_, *P*_6_]. Traversing in the opposite direction towards the other pivot *u*_6_ gives *V*_ℬ_ (*P*_1_, *P*_6_) = [*P*_1_, *P*_6_, *P*_5_, *P*_4_, *P*_3_, *P*_2_]. Now consider the relation in *L*_ℬ_ (*P*_*h*_, *P*_*p*_) for two distinct blocks *P*_*s*_, *P*_*t*_ ∈ *Part*(ℬ) that are neither the hybrid *P*_*h*_ nor the focal pivot *P*_*p*_.

#### Lemma 3.

*Let* 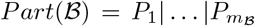 *be a partition induced by a non-trivial blob* ℬ *in a binary, semi-directed, level-1 network N* ^−^. *Let P*_*h*_ *be the hybrid block and P*_*p*_ *be a pivot block of Part*(ℬ). *Then, for all pairs of blocks P*_*s*_, *P*_*t*_ ∈ *Part*(ℬ) *such that s* ≠ *t* ≠ *h* ≠ *p, P*_*s*_ *< P*_*t*_ *is in L*_ℬ_ (*P*_*h*_, *P*_*p*_) *if and only if for all* 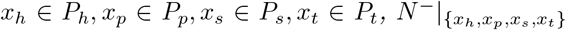 *displays quartet x*_*t*_, *x*_*h*_|*x*_*p*_, *x*_*s*_; *otherwise, P*_*t*_ *< P*_*s*_ *is in L*_ℬ_(*P*_*h*_, *P*_*p*_).

The proof of Lemma 3 is in the Appendix A. The idea is that one of the two displayed quartets will put the hybrid and the focal pivot together and the other will separate the hybrid and the focal pivot, revealing the linearized order of the non-pivot, non-hybrid blocks *P*_*s*_, *P*_*t*_. As the lowest of the three qCFs corresponds to the non-displayed quartet by Proposition 11 in [7], the ordering of *P*_*s*_, *P*_*t*_ can be determined by comparing the qCF values of the two quartets that separate the pivot and the hybrid (Algorithm 3 in Appendix A). This approach is closely related to the identifiability result for circular orders; however, the innovation behind NetCS is leveraging a pivot to linearize the order and then recovering it in an efficient manner via sorting (Algorithm 4 in Appendix A).

#### Theorem 3.

*Assume the same settings as Lemma 1. Let v*_ℬ_ *be some vertex in the TOB for N* ^−^ *with degree m*_ℬ_ ≥ 5. *Then, for every ϵ >* 0, *Algorithm 4 applied to Part*(*v*_ℬ_), *with hybrid block P*_*h*_ *and pivot block P*_*p*_, *and* 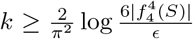 *gene trees returns the (correct) circular order of Part*(*v*_B_ ) *with probability at least* 1 − *ϵ. The total time complexity of applying Algorithm 4 to all v*_ℬ_ *is O*(*n* log *nk*) *in fast mode and O*(*n*^2^ log (*n*)*k*) *otherwise, where n is the number of taxa*.

*Proof*. By Lemma 3, Algorithm 3 returns the correct relation in *L*_ℬ_(*P*_*h*_, *P*_*p*_) for all pairs *P*_*s*_, *P*_*t*_ ∈ *Part*(*v*_ℬ_) such that *s*≠ *t*≠ *h*≠ *p* provided the lowest qCF corresponds to the non-displayed quartet for all 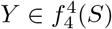 such that *N* ^−^|_*Y*_ contains a 4-blob. Following the same argument in the proof of Theorem 2, this occurs with probability at least 1 − *ϵ* when the number *k* of gene trees is at least 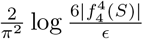. And, when the correct relations are returned, the (correct) circular order of *Part*(ℬ) is returned by merge sort (Algorithm 3–4).

Now consider time complexity. Merge sort makes *O*(*m*_ℬ_ log *m*_ℬ_) pairwise block comparisons. Each comparison requires qCF computation, which is *O*(*k*) time in fast mode or *O*(*nk*) time otherwise, as previously discussed for pivot scan. Repeating for all *v*_ℬ_ gives us our result. Circular sorting makes *O*(*n* log *n*) quarnet queries when running in fast mode.

### 3.4 Theoretical and Practical Considerations

Combining Theorems 1, 2, and 3 gives our final result:

#### Corollary 1.

*NetCS is statistically consistent under the NMSC and has time complexity O*(*n* log (*n*)*k*) *in fast mode or O*(*n*^2^ log (*n*)*k*) *otherwise for k genes and n species. In fast mode, NetCS computes qCFs for O*(*n* log *n*) *4-taxon subsets, referred to as quarnet queries*.

NetCS’s fast mode is for the regime where the error in all qCFs is below the tolerance threshold *π/*2, so every quarnet can be reconstructed with perfect accuracy (i.e., we have an oracle for quarnets). Only *O*(*n*) quarnets are accessed for hybrid voting and pivot scan; this increases to *O*(*n* log *n*) quarnets for circular sorting; thus, NetCS matches the lower bound of quarnet complexity recently established for semi-directed, level-1 network reconstruction (Proposition 6 in [13]). Moreover, when given direct access to quarnets, rather than inferring them from qCFs, NetCS has time complexity *O*(*n* log *n*). This improves upon the time complexity of full semi-directed level-1 network reconstruction from quarnets, which can be can be done in *O*(*n*^2^) using the taxon-addition-like algorithm recently presented by Frohn *et al*. (2025) (Theorem 13 in [13]). Like fast mode, the results in [13] are primarily of theoretical interest given the assumption of a perfect oracle (and there is no implementation of their algorithm to our knowledge).

Although our analysis primarily focused on fast mode, we initially designed the slower version NetCS, with the goal of increasing robustness to noisy qCFs, and only later defined the fast version to enable comparisons to [13]. Beyond the extra computation previously mentioned, our implementation of NetCS also runs circular sorting taking each block in *Part*(*v*_ℬ_), excluding the hybrid, as a candidate pivot, with the goal of increasing robustness to noisy qCFs. to increase the robustness. An error boolean is returned for each pairwise block comparison made during sorting with Algorithm 3, indicating whether the quartet with the hybrid and candidate pivot as siblings had the lowest qCF. This scenario should never occur if the candidate is the correct pivot by Theorem 2. Thus, NetCS reconstructs the blob taking the circular order with the lowest total error score, breaking ties to favor the candidate pivots originally returned from pivot scan and otherwise breaking ties arbitrarily. These additional steps do not impact statistical consistency because both pivots returned by the pivot scan will have score 0, with probability going to 1 as the number of gene trees goes to infinity, at which point, no other (incorrect) candidate can achieve a lower total error score. However, testing all candidate pivot blocks increases the total time complexity of NetCS to *O*(*n*^3^ log (*n*)*k*). As we will show, even with this additional computation, NetCS is still fast in practice likely because it avoids accessing qCFs for all subsets of four taxa. Precomputation of all qCFs not only has an *O*(*n*^4^*k*) time cost but also an Ω(*n*^4^) storage cost, which can lead to large constant factors in qCF access from the memory hierarchy (e.g., cache misses during either qCF precomputation or blob reconstruction).

NetCS is not without practical limitations, including those inherited from employing the same algorithmic framework of NANUQ+. First, an estimated TOB is required as input, which may contain varying levels of error. Second, all blobs are reconstructed independently, which can lead to an incompatible set of inferred hybrids (i.e., no valid semi-directed network exists that realizes all inferred hybrids simultaneously [18]). To address this issue, our implementation of NetCS processes blobs in decreasing order of degree and excludes blocks that conflict with previously inferred hybrids during the voting procedure. If all blocks are excluded, NetCS does not attempt to reconstruct that blob and thus fails to reconstruct the network. Likewise, NANUQ+ can fail to produce a valid output network due to conflicts among inferred hybrids.

### 3.5 Experimental Evaluation

To study the empirical performance of NetCS, we conducted an evaluation study using data sets simulated under the NMSC, so the true level-1 network and its tree of blobs were known. Details, including software commands and version numbers, are given in Appendix B.

#### Experiment 1

The goal of experiment 1 was to evaluate the accuracy of level-1 blob reconstruction with NetCS. Data sets [20] used for this evaluation were previously simulated by [10] with 12 model conditions corresponding to three numbers of taxa (50, 100, and 200) and four branch length scaling factors (2.0, 1.0, 0.5, 0.25). The scaling factors varied the level of gene tree discordance due to ILS (note that the mean discordance levels were 45%, 65%, 80%, and 90%, respectively, based on simulated level-0 networks). Twenty replicate networks were available per model condition. For each replicate, 1000 gene trees were simulated under the NMSC with independent inheritance. The number of (true) gene trees given as input to NetCS was varied from 100, 500, and 1000, yielding 36 conditions in total.

NetCS was given the true TOB as input so that hybrid detection and circular order reconstruction could be evaluated for blobs with degree at least 5. **Hybrid error** was defined as the proportion of blobs for which the predicted hybrid block differed from the true hybrid block. **Circular order error** was quantified using the Kendall tau distance. For two linear permutations *σ*_1_, *σ*_2_ on the set *B* = {1, 2, …, *m*_ℬ_}, the Kendall tau distance *D*_*kτ*_ (*σ*_1_, *σ*_2_) is the fraction of unordered pairs (*i, j*) ∈ *B* whose relative ordering differs between *σ*_1_ and *σ*_2_ [9]. We extended this notion to circular orders by considering all linearizations obtained by cutting the circle at each position as well as its reversal. Let *Σ*(*ζ*) denote the set of all linearizations of circular order *ζ*, and let *Σ*(*ζ*)^*R*^ denote their reversals. Then, the Kendall tau distance between two circular orders *ζ*_1_, *ζ*_2_ is

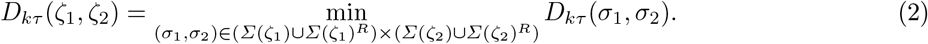

We reported hybrid error and circular order error averaged over all blobs of degree at least 5 in the true tree of blobs for each replicate network.

#### Experiment 2

The goal of experiment 2 was to benchmark NetCS against NANUQ+ in terms of blob reconstruction error as well as computational efficiency. This evaluation used the performance metrics and data sets as experiment (1) except that true gene trees were replaced with gene tree estimated from simulated DNA sequences using IQ-TREE model selection [26, 21] (sometimes gene tree estimation from simulated DNA sequences failed but the total number of estimated gene trees was always greater than 900). Mean gene tree estimation error, quantified in terms of (normalized) Robinson-Foulds error [31], ranged from 40% to 70%. Both NetCS and NANUQ+ were given the true TOB as input so that reconstructed blobs with degree 5 or more could be compared. Computational efficiency was evaluated as wallclock time (24-hour limit) and peak resident set size (peak RSS; 256 GB limit) (note that the reported time and memory usage does not include TOB reconstruction, as the true TOB was given as input). Both methods were run on compute nodes with the same architecture (AMD EPYC 7313) when collecting these performance metrics. NANUQ+ was given access to 8 threads for qCF precomputation.

#### Experiment 3

The goal of the third experiment was to evaluate NetCS for end-to-end semi-directed level-1 network reconstruction in comparison with NANUQ+ and CAMUS. Data sets used for this evaluation were previously simulated for the study presenting CAMUS [39]. The CAMUS data sets had four model conditions with 51, 101, 151, and 201 taxa, including a single taxon outgroup (note: CAMUS could not be run in the previous experiments because the networks did not have a single taxon outgroup, which led to issues correctly rooting CAMUS’ estimated base tree). Replicate networks with blobs of degree 4 were excluded, as the hybrid is not identifiable for those blobs; this left 47, 45, 43, and 49 replicates (out of 50) for the 51-, 101-, 151-, and 201-taxon conditions, respectively. Each replicate had 1000 gene trees. Mean gene tree estimation error was ∼ 20%; ILS level was not reported by [38].

Since experiment 3 was based on data sets from the study presenting CAMUS [38], CAMUS was run with the same default and recommended settings. The base tree for CAMUS was estimated with ASTRAL-IV [42] and rooted at the outgroup taxon. Unlike the other methods, CAMUS returns a *collection* of (binary) *directed* level-1 networks, one after each edge addition; we used the directed network with the same number of edge additions as the number of non-trivial blobs (i.e., hybrid vertices) as the true network, with the goal of giving CAMUS an advantage. For running NetCS and NANUQ+, the ASTRAL-IV tree was used to estimate a TOB via edge contractions following the approach of [10]; see Appendix B for details. NetCS and NANUQ+ were run twice: once given the estimated TOB as input and once given the true TOB as input.

All methods were evaluated based on hybrid ancestry error and network topology error. A species has **hybrid ancestry** if it is a hybrid or is the descendant of one or more hybrids. False positives (FPs) correspond to the estimated network predicting a species has hybrid ancestry when it does not according to the true network; false negatives (FNs) are the reverse. FPR is the number of FPs divided by the number of species with hybrid ancestry according to the estimated network. FNR is the number of FNs divided by the number of species with hybrid ancestry according to the true network. Balanced error rate, defined as the arithmetic mean of the FPR and FNR, was reported for all methods. Network topology error was evaluated as the **normalized hardwired cluster distance (nHWCD)** [19]. Each edge in a directed network induces a hardwired cluster of all the taxon descendants of the edge. The hardwired clusters induced by all edges of the network form the hardwired cluster system. The HWCD is the symmetric difference of the hardware cluster systems of two directed networks. When comparing level-1 networks, HWCD is a distance [25, 6, 8]. The extension to semi-directed networks takes the minimum value from all possible ways to “root” the two networks [18]. The normalized version divides HWCD by the total number of edges in both networks minus the total number of leaves in both networks. We used the implementation of nHWCD in PhyloNetworks [35], treating all networks as semi-directed.

## 4 Experimental Results

We now present the results of the three experiments.

### Experiment 1

The goal of experiment 1 was to evaluate how model condition impacted level-1 blob reconstruction with NetCS (Table 2, Fig. 2). As the number of gene trees and the branch scale factor increased, error decreased. For example, the average Kendall tau distance dropped from 0.14 to 0.00 going from the hardest model condition (0.25× branch scaling factor and 100 true gene trees) to the easiest (2.0× branch scaling factor and 1000 true gene trees), both with 50 taxa. These trends are expected as larger numbers of true gene trees correspond to lower error in observed qCFs and larger branch length scaling factors correspond to lower ILS and thus higher error tolerance *π*. Similar trends were observed for model conditions with larger numbers of taxa and hybrid error. Average hybrid error was generally low at 1000 gene trees, ranging from 0.12 to 0.00 (see Table 2 for details). NetCS did not have any failures in experiment 1.

**Fig. 2:**
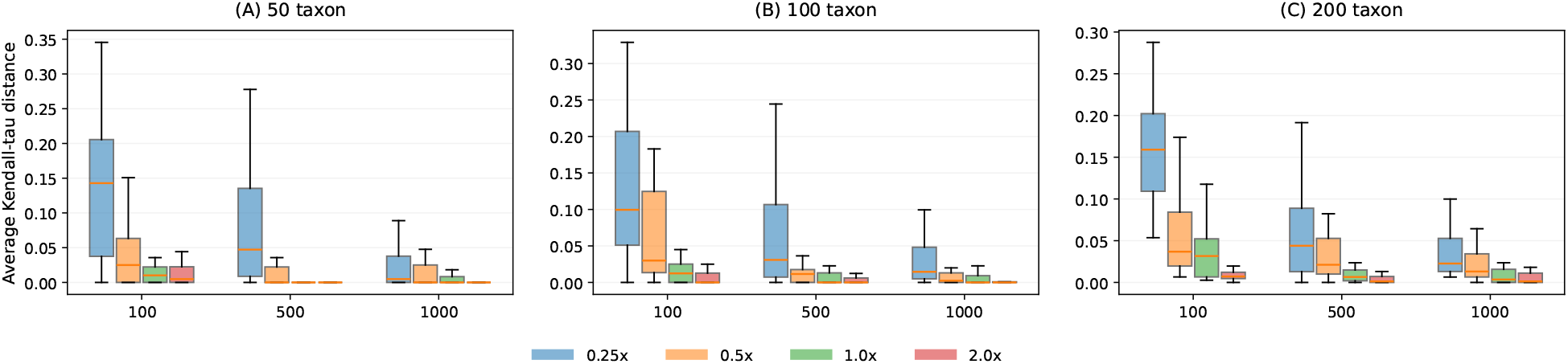
Experiment 1: Average Kendall-tau distance. Three subfigures (A), (B),(C) correspond to data sets with 50 taxa, 100 taxa, and 200 taxa, respectively. The *x*-axis is the number of gene trees. Each colored box represents a different branch scaling factor in order of decreasing ILS. The *y*-axis is error in circular order reconstruction (i.e., Kendall-tau distance, averaged across all blobs in replicate). Orange lines represent medians. Outliers are not shown.

### Experiment 2

The goal of experiment 2 was to compare NetCS against NANUQ+ in terms of level-1 blob reconstruction error and computational efficiency (Table 1, Fig. 6). Again, the percent hybrids correctly predicted increased with decreasing ILS level, with NetCS making the correct call for 79%, 83%, 88%, and 95% of blobs in 50-taxon data sets, respectively. Net-CS outperformed NANUQ+ in hybrid detection on 5 model conditions, the reverse occurred on 1 model condition, and the two methods tied in the remaining 2 model conditions. Error in circular order reconstruction also decreased with increasing ILS level, with NetCS and NANUQ+ tied, each outperforming the other on 4 conditions. The difference between methods for both tasks was small, suggesting they achieved similar accuracy.

**Table 1:**
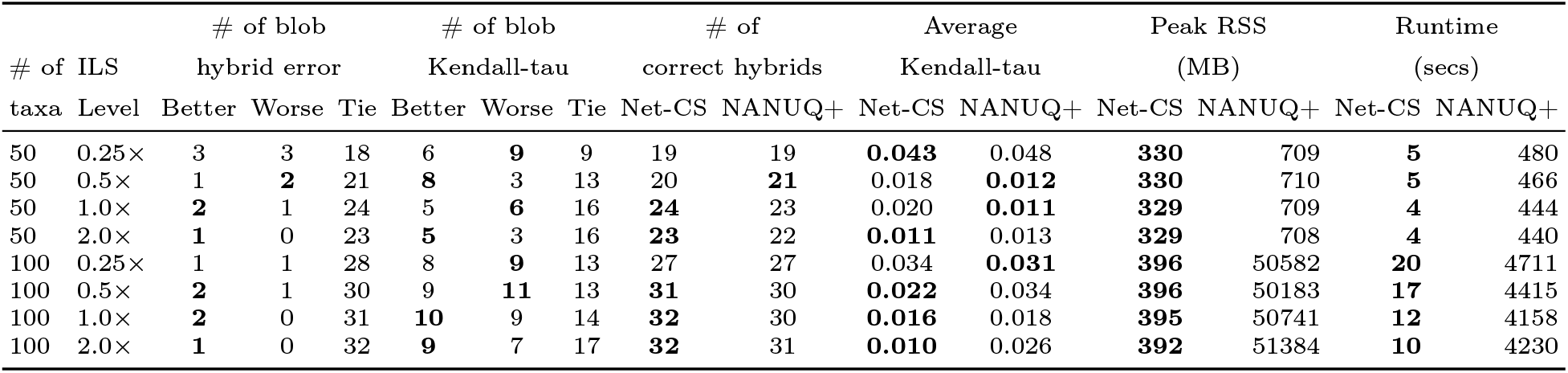
Experiment 2: The table shows the number of blobs, across all replicate networks for the same model condition, in which NetCS is better / worse / tied with NANUQ+ in terms of hybrid error and circular order error (Kendall tau distance). The Kendall-tau distance is the average across all blobs in a replicate, which is then averaged across all replicates. Runtime and peak RSS values are averages across all replicates per model condition. Results are not shown for 200 taxa, as NANUQ+ did not complete within the maximum wallclock time of 24 hours or exceeded the memory limit of 256GB. Both methods were given the same estimated gene trees as input (∼1000).

|  |  | # of blob |  |  | # of blob |  |  | # of |  | Average |  | Peak RSS |  | Runtime |  |
| --- | --- | --- | --- | --- | --- | --- | --- | --- | --- | --- | --- | --- | --- | --- | --- |
| # of ILS |  | hybrid error |  |  | Kendall-tau |  |  | correct hybrids |  | Kendall-tau |  | (MB) |  | (secs) |  |
| taxa | Level | Better | Worse | Tie | Better | Worse | Tie | Net-CS | NANUQ+ | Net-CS | NANUQ+ | Net-CS | NANUQ+ | Net-CS | NANUQ+ |
| 50 | 0.25× | 3 | 3 | 18 | 6 | **9** | 9 | 19 | 19 | **0.043** | 0.048 | **330** | 709 | **5** | 480 |
| 50 | 0.5× | 1 | **2** | 21 | **8** | 3 | 13 | 20 | **21** | 0.018 | **0.012** | **330** | 710 | **5** | 466 |
| 50 | 1.0× | **2** | 1 | 24 | **5** | **6** | 16 | **24** | 23 | 0.020 | **0.011** | **329** | 709 | **4** | 444 |
| 50 | 2.0× | **1** | 0 | 23 | **5** | 3 | 16 | **23** | 22 | **0.011** | 0.013 | **329** | 708 | **4** | 440 |
| 100 | 0.25× | 1 | 1 | 28 | 8 | **9** | 13 | 27 | 27 | 0.034 | **0.031** | **396** | 50582 | **20** | 4711 |
| 100 | 0.5× | **2** | 1 | 30 | 9 | **11** | 13 | **31** | 30 | **0.022** | 0.034 | **396** | 50183 | **17** | 4415 |
| 100 | 1.0× | **2** | 0 | 31 | **10** | 9 | 14 | **32** | 30 | **0.016** | 0.018 | **395** | 50741 | **12** | 4158 |
| 100 | 2.0× | **1** | 0 | 32 | **9** | 7 | 17 | **32** | 31 | **0.010** | 0.026 | **392** | 51384 | **10** | 4230 |

**Table 2:** Experiment 1. Each row shows results for particular model condition defined by the number of taxa and branch scaling factor, which in turn varies ILS level. Evaluation metrics are averaged over all replicates for the same model condition. Columns indicate number of gene trees, varying from 100, 500, to 1000. Hybrid error is the fraction of incorrect hybrid across all blobs in a network. Kendall-tau distance is averaged across all blobs in a network.

| # of scale<br>taxa factor |  | Hybrid<br>error |  |  | Kendall-tau<br>distance |  |  | Peak RSS<br>(MB) |  |  | Runtime<br>(secs) |  |  |
| --- | --- | --- | --- | --- | --- | --- | --- | --- | --- | --- | --- | --- | --- |
|  |  | 100 | 500 | 1000 | 100 | 500 | 1000 | 100 | 500 | 1000 | 100 | 500 | 1000 |
| 50 | 0.25× | 0.708 | 0.392 | 0.117 | 0.144 | 0.082 | 0.032 | 273 | 299 | 331 | 4 | 5 | 5 |
| 50 | 0.5× | 0.308 | 0.092 | 0.100 | 0.050 | 0.015 | 0.012 | 273 | 299 | 330 | 7 | 4 | 6 |
| 50 | 1.0× | 0.142 | 0.100 | 0.108 | 0.017 | 0.003 | 0.020 | 274 | 298 | 329 | 4 | 3 | 4 |
| 50 | 2.0× | 0.100 | 0.050 | 0.000 | 0.022 | 0.004 | 0.003 | 274 | 298 | 328 | 4 | 3 | 4 |
| 100 | 0.25× | 0.525 | 0.142 | 0.108 | 0.136 | 0.058 | 0.033 | 282 | 333 | 395 | 12 | 11 | 17 |
| 100 | 0.5× | 0.100 | 0.025 | 0.000 | 0.070 | 0.031 | 0.016 | 283 | 332 | 395 | 8 | 11 | 13 |
| 100 | 1.0× | 0.042 | 0.025 | 0.025 | 0.028 | 0.020 | 0.011 | 283 | 332 | 395 | 7 | 7 | 10 |
| 100 | 2.0× | 0.017 | 0.000 | 0.000 | 0.008 | 0.004 | 0.005 | 285 | 334 | 395 | 5 | 8 | 9 |
| 200 | 0.25× | 0.650 | 0.042 | 0.054 | 0.168 | 0.057 | 0.034 | 300 | 398 | 524 | 40 | 40 | 65 |
| 200 | 0.5× | 0.096 | 0.037 | 0.025 | 0.054 | 0.038 | 0.025 | 299 | 399 | 525 | 25 | 31 | 55 |
| 200 | 1.0× | 0.062 | 0.025 | 0.050 | 0.045 | 0.019 | 0.019 | 302 | 401 | 526 | 26 | 25 | 49 |
| 200 | 2.0× | 0.013 | 0.025 | 0.025 | 0.015 | 0.010 | 0.011 | 303 | 403 | 528 | 19 | 21 | 46 |

In contrast, NetCS and NANUQ+ differed substantially in terms of their computational efficiency. On the 50-taxon data sets, Net-CS completed in under 5 seconds on average using less than 0.33GB memory, whereas NANUQ^+^ required ∼7–8 minutes on average using over 0.7GB memory. The performance gap increased for the 100-taxon data sets. Net-CS completed in 20 seconds on average using less than 0.4GB memory, whereas NANUQ^+^ required more than one hour on average using over 50GB memory. On the 200-taxon data sets, NANUQ^+^ either exceeded the 24-hour wall-clock limit or the 256GB memory limit, while NetCS completed within 15–215 seconds using ∼0.5GB memory. Note that NetCS did not have any failures for experiment 2, but NANUQ+ failed to return a valid network for one 50-taxon network (id #1207) for branch scaling factors (0.25×, 0.5×, 2.0×) and one 100-taxon network (id #10032) for one branch scaling factor (0.25×); these replicate data sets were excluded to ensure fair error and runtime comparisons.

### Experiment 3

The goal of the final experiment was to benchmark NetCS as part of an end-to-end pipeline for network reconstruction against NANUQ+ and CAMUS. NANUQ+ (given estimated TOBs) failed to return a valid network for 2 replicates with 50 taxa and 2 with 100 taxa, and it did not complete on 150- and 200-taxon data sets within the maximum wallclock time of 24 hours or exceeded the memory limit of 256GB. NetCS (given estimated TOBs) failed to return a valid network for 1 replicate with 150 taxa, respectively. These replicates were excluded to ensure fair comparison.

For network topology error, CAMUS, NetCS (with estimated TOB), and NANUQ+ (with estimated TOB) produced networks with nHWCD between 0.05–0.15 (Fig. 3). Although CAMUS slightly outperformed NetCS on 50 taxa (median nHWCD: 0.08 vs. 0.10), NetCS slightly outperformed CAMUS on 100 taxa (median nHWCD: 0.06 vs. 0.08). The two methods had similar performance on 150 and 200 taxa. NetCS slightly outperformed NANUQ+ in terms of topological error: median nHWCD of 0.10 vs. 0.13 and 0.06 vs. 0.08 for 50 and 100 taxa, respectively.

**Fig. 3:**
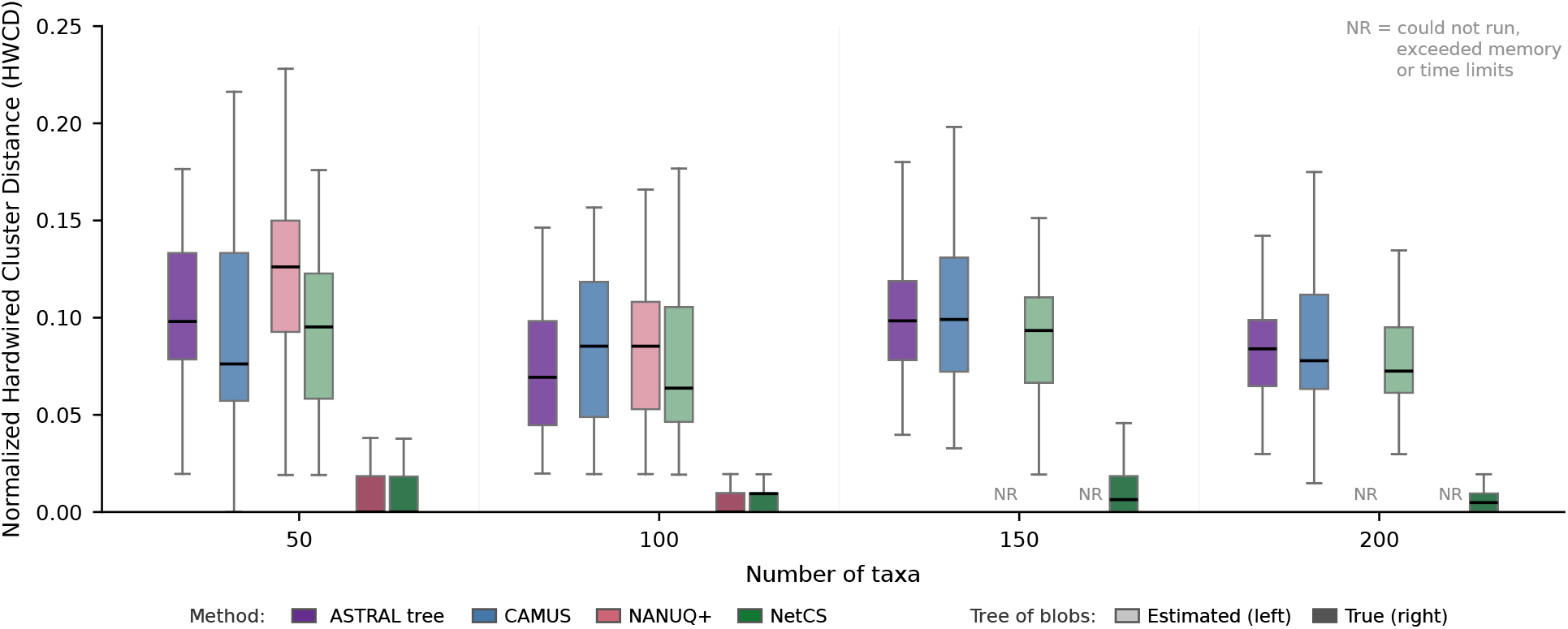
Experiment 3: Normalized HWCD. The *y*-axis is network topological error. The *x*-axis is the number of taxa. Each boxplot is the results from applying a method on replicate data sets with the specified number of taxa. Bold black lines represent medians; outliers are not shown. NANQU^+^ failed to run on 150- and 200-taxon datasets, annotated as “NR”.

Overall, the topological error of all methods was typically close to that of the ASTRAL-IV tree. Providing NetCS and NANUQ+ with the true TOBs, instead of an estimated TOB reconstructed from the ASTRAL-IV tree, dropped nHWCD to near 0 for all numbers of taxa. Unlike NetCS and NANUQ+, CAMUS adds edges to the ASTRAL-IV tree. The nHWCD of the resulting network was better/worse than the ASTRAL-IV tree for 27/18, 19/24, 22/20, and 27/22 replicates for the 51-, 101-, 151-, and 201-taxon model conditions; thus, the edge additions made by CAMUS decreased nHWCD slightly more often then they increased it.

For hybrid ancestry error, all methods exhibited poor performance (Fig. 4). CAMUS had the highest variability, sometimes achieving balanced error close to 0 and other times close to 1; however, its median balanced error rate was greater than 0.5 on all model conditions with more than 50 taxa. NetCS outperformed CAMUS and NANUQ+ in terms of hybrid ancestry predictions on these conditions, although its balanced error rate was still high. Providing NetCS and NANUQ+ with the true TOB dropped the balanced error rate to near 0 for all numbers of taxa. Note the (balanced) topological error rate of the estimated TOBs was 0.06–0.08 on average, with FPR consistently higher than FNR (Table 3 in the Appendix C).

**Table 3:** Experiment 3: TOB topological error. Number of taxa in model condition for CAMUS data sets. Number of replicates remaining after excluding those whose ground truth network contains one or more 4-blob. Error between the true and estimated TOB are reported as FNR and FPR, defined in Figure 5 caption. Error values are mean ± standard deviation across replicates.

| Taxa # | of Replicates | FNR | FPR | (FNR+FPR)/2 |
| --- | --- | --- | --- | --- |
| 50 | 47 | $0.039 \pm 0.029$ | $0.120 \pm 0.053$ | $0.079 \pm 0.033$ |
| 100 | 45 | $0.038 \pm 0.017$ | $0.078 \pm 0.040$ | $0.058 \pm 0.024$ |
| 150 | 43 | $0.039 \pm 0.018$ | $0.088 \pm 0.034$ | $0.064 \pm 0.022$ |
| 200 | 49 | $0.040 \pm 0.014$ | $0.077 \pm 0.022$ | $0.059 \pm 0.015$ |

**Fig. 4:**
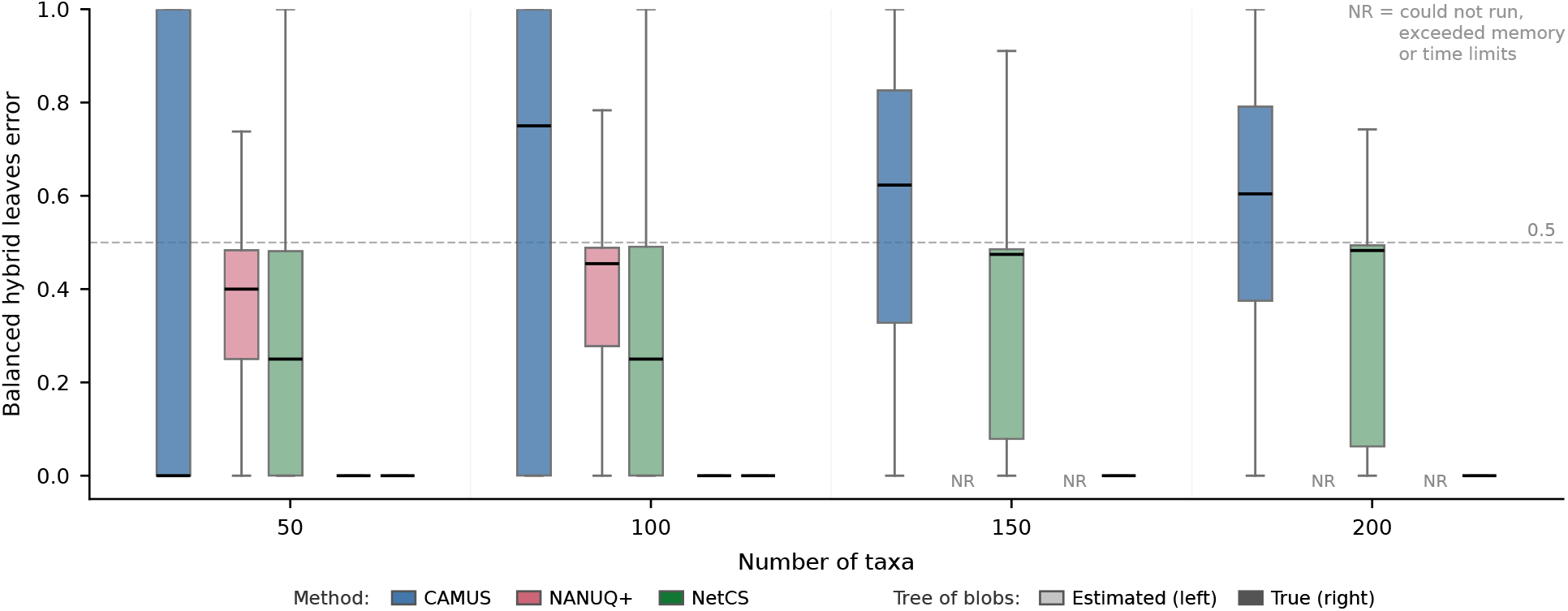
Experiment 3: Hybrid ancestry balanced error rate. The *y*-axis is the balanced hybrid ancestry error rate. The *x*-axis is the number of taxa. Each boxplot is the results from applying a method on replicate data sets with the specified number of taxa. Bold black lines represent medians; outliers are not shown. NANQU^+^ failed to run on 150- and 200-taxon datasets, annotated as “NR”.

## 5 Conclusions

In this paper, we studied the problem of reconstructing level-1 blobs under the NMSC model when the tree of blobs is known. This work was motivated by recent advances in tree of blobs reconstruction [3, 17, 13, 10, 30] and by the two-step design of NANUQ+ for reconstructing semi-directed level-1 networks. Our method, NetCS, combines majority voting to identify the hybrid vertex with sorting to recover the circular order. These primitives closely reflect prior identifiability results for semi-directed level-1 networks [34, 7, 1], although, to our knowledge, they had not previously been operationalized in a fast, statistically consistent method.

A key contribution of NetCS is that it avoids the precomputation of all qCFs by (1) repurposing the 3f1a algorithm from our recent study [10] and (2) applying an efficient sorting algorithm. This gives NetCS time complexity *O*(*n* log (*n*)*k*) and only *O*(*n* log *n*) quarnet queries, matching the lower bound of [13]. These results indicate that consistent reconstruction of level-1 blobs under the NMSC is easy, at least from a theoretical perspective.

Notably, our implementation of NetCS performs additional computation to improve robustness to noisy qCF estimates, leading to *O*(*n*^3^ log (*n*)*k*) time complexity. Nevertheless, by avoiding the quartic cost of qCF precomputation, NetCS enables analyses of data sets with 200 taxa and 1000 genes in only a few minutes, substantially improving upon NANUQ+ in both runtime and memory usage. Despite its reduced access to qCFs, NetCS was still highly competitive with NANUQ+ in terms of accuracy. However, when supplied with an estimated tree of blobs, the accuracy of both methods deteriorated. This finding suggests that the primary practical challenge may lie in fast and accurate tree of blobs reconstruction.

Direct network reconstruction circumvents this issue, motivating us to benchmark NetCS and NANUQ+ against the recently proposed method CAMUS (we did not evaluate SNaQ as it was less accurate than CAMUS on the same data sets). CAMUS, NetCS, and NANUQ+ had comparable network topology error. CAMUS, in particular, struggled to accurately determine whether species evolved through hybridization, a question of importance in biological studies. NetCS (given the estimated tree of blobs) performed better than CAMUS with respect to this criterion for data sets with 100 or more taxa, although its performance was still poor.

Collectively, these findings highlight the need for improved methods for level-1 network reconstruction. Given the computational efficiency and strong performance of NetCS when provided with the true tree of blobs, a promising direction for future research is to combine NetCS with improved methods for tree of blobs reconstruction. The algorithmic techniques underlying NetCS also could be explored in the context of direct network reconstruction or other approaches for level-1 blob reconstruction. For example, reducing the number of 4-taxon subsets sampled in NANUQ+ or replacing collections of 4-taxon subsets with averages. Such techniques could also be relevant for applications of the recent method Squirrel [16] to gene tree data. Squirrel is similar to NANUQ+, in that both are based on quartet distances, but was introduced in the context of level-1 blob reconstruction from DNA sequences.

## A Supplemental Methods

### Lemma 2.

*Let Part* 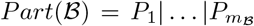 *be a partition induced by a non-trivial blob* ℬ *in a binary, semi-directed, level-1 network N* ^−^, *and let P*_*h*_ *be the hybrid block of Part*(ℬ). *Then, P*_*p*_ *is a pivot block of Part*(ℬ) *if and only if for all s, t* ∈ {1, …, *m*_ℬ_} *such that s* ≠ *t*≠ *p* ≠ *h and all x*_*h*_ ∈ *P*_*h*_, *x*_*p*_ ∈ *P*_*p*_, *x*_*s*_ ∈ *P*_*s*_, *x*_*t*_ ∈ *P*_*t*_, 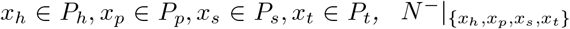 *displays quartet x*_*h*_, *x*_*p*_|*x*_*s*_, *x*_*t*_.

*Proof*. Because *x*_*h*_, *x*_*p*_, *x*_*s*_, *x*_*t*_ are drawn from distinct blocks of *Part*(B) and *x*_*h*_, in particular, is drawn from the hybrid block, 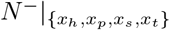 contains a 4-blob by Lemma 5 in [10]. Let *u*_*j*_ denote the vertex in B such that the deletion of the one cut edge incident to *u*_*j*_ induces the bipartition: *P*_*j*_|*S* \ *P*_*j*_ for all *j* ∈ {1, …, *m*_*ℬ*_}.

⇒ Assume that *P*_*p*_ is a pivot block of *Part*(ℬ). By definition, there exists a reticulation edge (*u*_*p*_, *u*_*h*_) in *N* ^−^. After deleting the partner reticulation edge of (*u*_*p*_, *u*_*h*_), any path of distinct edges between *x*_*h*_ ∈ *P*_*h*_ and *x*_*p*_ ∈ *P*_*p*_ in 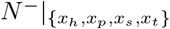 contains edge (*u*_*p*_, *u*_*h*_) but no other edges in B. In contrast, any path of distinct edges between *x*_*h*_ and *x*_*s*_ ∈ *P*_*s*_ or *x*_*t*_ ∈ *P*_*t*_ in 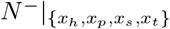 contains edge (*u*_*p*_, *u*_*h*_) as well as the (non-reticulation) edge in that is incident to (*u*_*p*_, *u*_*h*_), which induces bipartition *x*_*h*_, *x*_*p*_ | *x*_*s*_, *x*_*t*_ giving us our result.

⇐ For the sake of obtaining a contradiction, assume the condition holds but *P*_*p*_ is not a pivot block. Then, there exists two pivot blocks *P*_*v*_, *P*_*w*_ where *w*≠ *v* ≠ *p*≠ *h*. For all *x*_*h*_ ∈ *P*_*h*_, *x*_*p*_ ∈ *P*_*p*_, *x*_*v*_ ∈ *P*_*v*_, *x*_*w*_ ∈ *P*_*w*_, 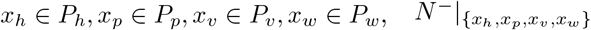 contains a 4-blob that displays two quartets: *x*_*h*_, *x*_*v*_|*x*_*w*_, *x*_*p*_ from deleting reticulation edge (*u*_*w*_, *u*_*h*_) and *x*_*h*_, *x*_*w*_|*x*_*v*_, *x*_*p*_ from deleting reticulation edge (*u*_*v*_, *u*_*h*_), as described in the forward direction of the proof. As 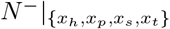 displays two quartets given our assumption that *N* ^−^ is level-1 and binary [7], quartet *x*_*h*_, *x*_*p*_|*x*_*v*_, *x*_*w*_ is not displayed, which is a contradiction.

### Algorithm 2

Pivot Scan Algorithm

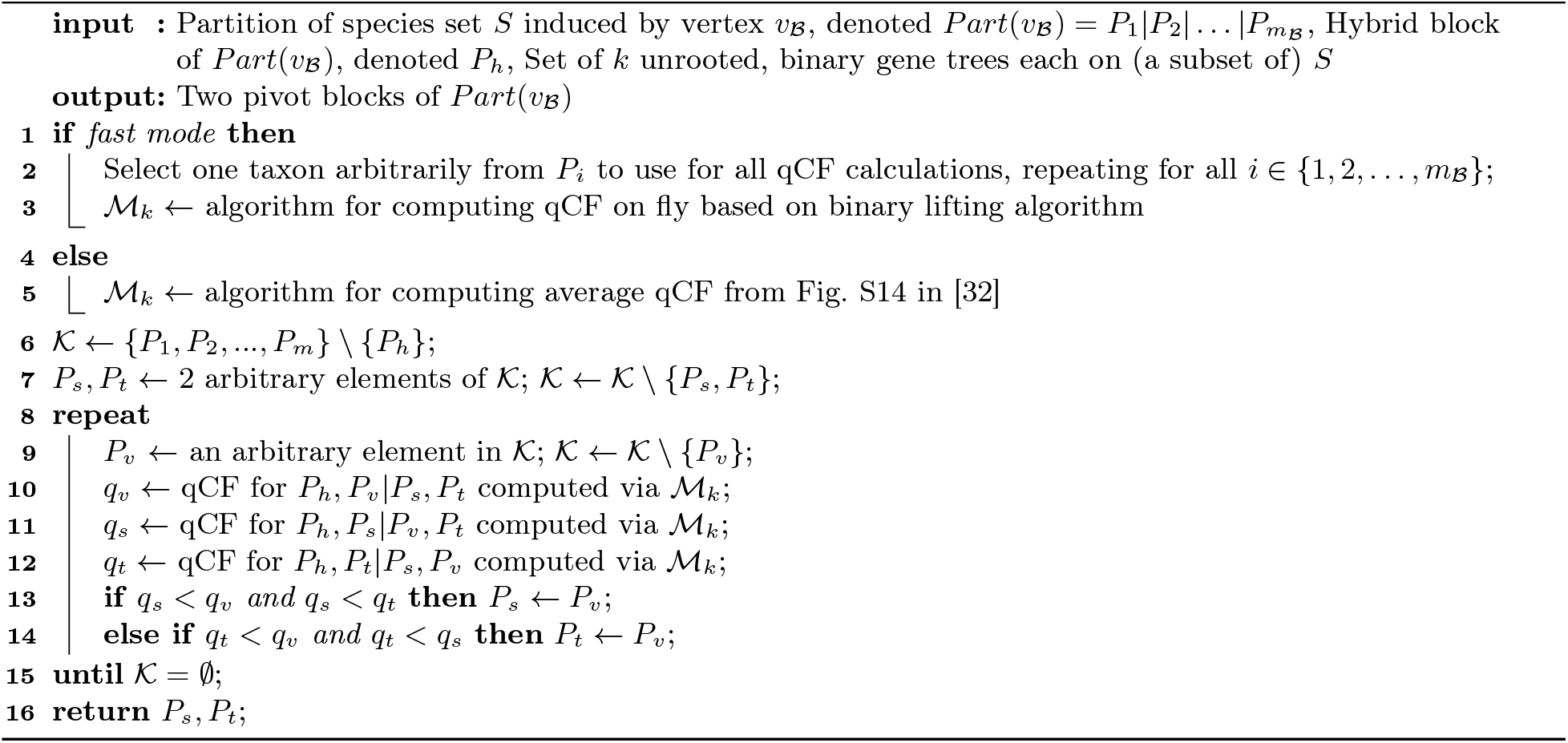

### Lemma 3.

*Let Part* 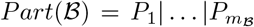 *be a partition induced by a non-trivial blob* ℬ *in a binary, semi-directed, level-1 network N* ^−^. *Let P*_*h*_ *be the hybrid block and P*_*p*_ *be a pivot block of Part*(ℬ). *Then, for any two blocks P*_*s*_, *P*_*t*_ ∈ *Part*(ℬ) *such that s t*≠ *h*≠ *p, P*_*s*_ *< P*_*t*_ *is in L*_*B*_(*P*_*h*_, *P*_*p*_) *if and only if for all x*_*h*_ ∈ *P*_*h*_, *x*_*p*_ ∈ *P*_*p*_, *x*_*s*_ ∈ *P*_*s*_, *x*_*t*_ ∈ *P*_*t*_, 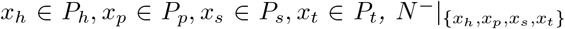 *displays quartet x*_*t*_, *x*_*h*_|*x*_*p*_, *x*_*s*_; *otherwise, P*_*t*_ *< P*_*s*_ *is in L*_ℬ_ (*P*_*h*_, *P*_*p*_).

*Proof*. Because *x*_*h*_, *x*_*p*_, *x*_*s*_, *x*_*t*_ are all drawn from distinct blocks of *Part*(B) and *x*_*h*_, in particular, is drawn from the hybrid block, 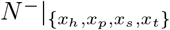 contains a 4-blob by Lemma 5 in [10]. Let *u*_*j*_ denote the vertex in ℬ such that the deletion of the one cut edge incident to *u*_*j*_ induces the bipartition: *P*_*j*_|*S* \ *P*_*j*_ for all *j* ∈ {1, …, *m*_*B*_}.

⇒ Assume *P*_*s*_ *< P*_*t*_ is in *L*_ℬ_ (*P*_*h*_, *P*_*p*_). Then, traversing ℬ from the hybrid vertex *u*_*h*_ toward pivot *u*_*p*_ encounters *u*_*s*_ before *u*_*t*_. It follows that the restriction of *N* ^−^ to {*x*_*h*_, *x*_*p*_, *x*_*s*_, *x*_*t*_} has two reticulation edges (*u*_*p*_, *u*_*h*_) and (*u*_*t*_, *u*_*h*_), and deleting reticulation edge (*u*_*p*_, *u*_*h*_) gives the displayed quartet *x*_*t*_, *x*_*h*_|*x*_*p*_, *x*_*s*_.

⇐ For the sake of obtaining a contradiction, assume the condition holds but *P*_*t*_ *< P*_*s*_ is in *L*_ℬ_ (*P*_*h*_, *P*_*p*_). The restriction of *N* ^−^ to {*x*_*h*_, *x*_*p*_, *x*_*s*_, *x*_*t*_} displays quartets *x*_*s*_, *x*_*h*_|*x*_*p*_, *x*_*t*_ and *x*_*h*_, *x*_*p*_|*x*_*s*_, *x*_*t*_ by the forward direction above and by Lemma 2, respectively. Only two quartets can be displayed given our assumption that *N* ^−^ is level-1 and binary [7]; thus, quartet *x*_*t*_, *x*_*h*_|*x*_*p*_, *x*_*s*_ is not displayed, which is a contradiction.

### Algorithm 3

Linearized Circular Order Comparison Algorithm

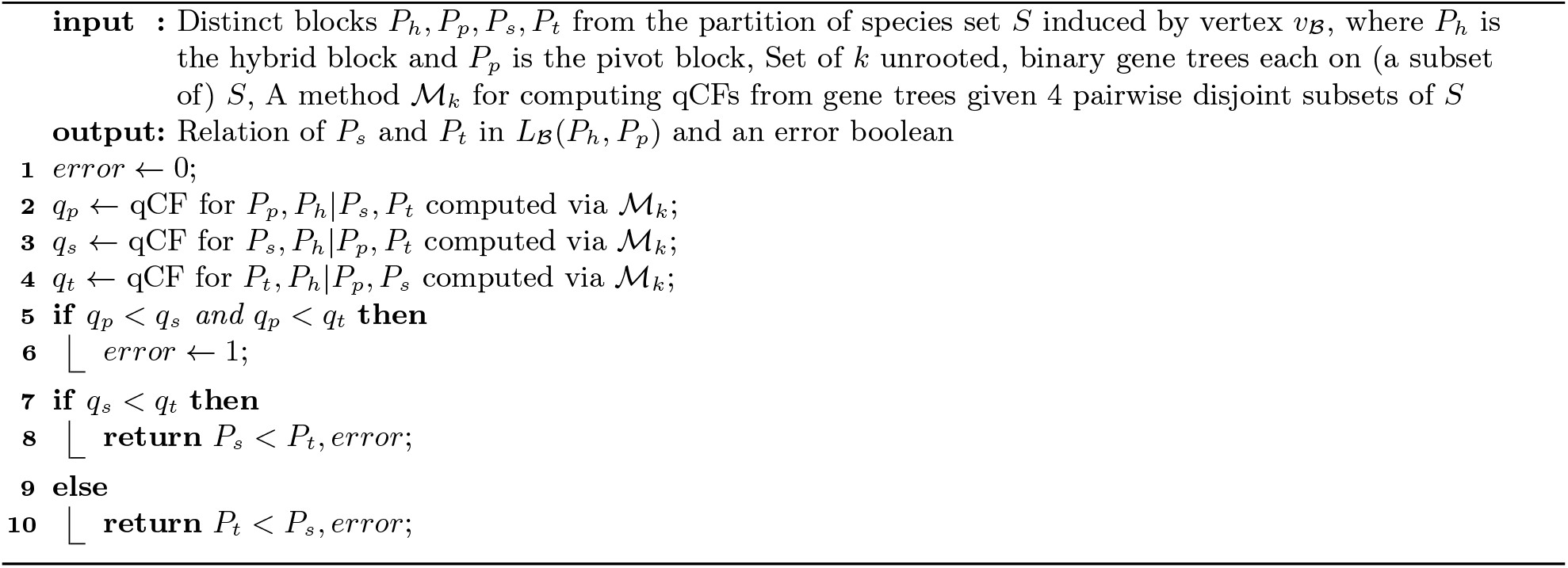

### Algorithm 4

Linearized Circular Sorting Algorithm

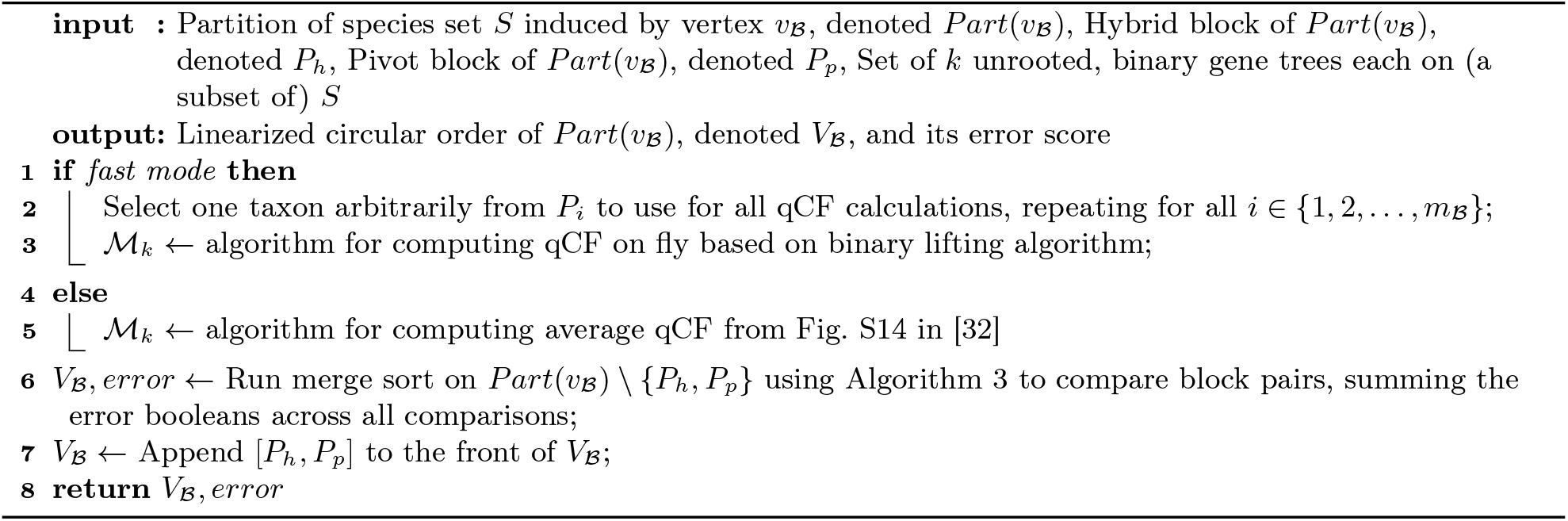

## B Software Commands

### B.1 NetCS Command

NetCS (available at https://github.com/molloy-lab/TREE-QMC; commit e577f8c for experiments 1 and 2 and commit eb3e140 for experiment 3) was run with command:

~~~
tree −qmc −−network −i [ input gene t r e e ] −−a t [ input TOB] −o [ ouput ]
~~~

### B.2 NANUQ+ Commands

NANUQ+, implemented in MSCQuartets v3.2 (https://cran.r-project.org/web/packages/MSCquartets/index.html), was run with commands:

~~~
l i b r a r y ( ape )
l i b r a r y ( MSCquartets )
g t t r e e <− read . t r e e ( “[ input gene t r e e s ]” )
QT <− q u a r te t Ta b l e P a r a l l e l ( g ttr e e, numCores = 8 )
RQT <− quartetTableToRQT (QT)
pT <− quartet Tree Test Ind (RQT, model=t3_model )
pT <− quartet Star Test Ind (pT)
ToB <− read . t r e e ( “[ input TOB]” )
ToB <− l a b e l In t N o d e s (ToB, p l o t=FALSE)
nets <− r e s o l v e L e v e l 1 (ToB, pT, alpha =[ alpha ], beta =[ beta ], p l o t=FALSE)
f inal_nwk <− nets [ [ 1 ] ] [ [ 1 ] ]
~~~

For experiments 1–2, NANUQ+ was given the true TOB and significance thresholds: *α* = 10^−7^, *β* = 0.99 based on hyperparameter tuning the study simulating the data sets [10]. For experiment 3, NANUQ+ was given a TOB estimated with TOB-QMC and significance thresholds: *α* = 10^−4^, *β* = 0.99, as described in Section B.4.

### B.3 ASTRAL-IV Command

ASTRAL-IV (v1.23.4.6), implemented in ASTER (https://github.com/chaoszhang/ASTER), was run with command:

~~~
a s t r a l 4 −t 32 −o [ output ] [ input gene t r e e s ]
~~~

The resulting tree was rooted at the outgroup using the scripts provided by the CAMUS study [38] with the following command:

python3 root−outgroup . py [ASTRAL t r e e ]

### B.4 TOB-QMC commands

TOB-QMC-3f1a (available at https://github.com/molloy-lab/TREE-QMC; commit e577f8c for experiments 1 and 2 and commit eb3e140 for experiment 3) was run with the command:

~~~
tree −qmc −−b l o b s e a r c h o n l y [ input ASTRAL t r e e ] \
−−s tore_pvalue −−3f 1 a −−o v e r r i d e \
−i [ input gene t r e e s ] \
−o [TOB refinement t r e e ]
~~~

to annotate the refinement tree computed with ASTRAL-IV with *p*-values using NANUQ+’s hypothesis test (T3 option). TOBs were constructed by contracting branches based on varying significance thresholds with the following command:

~~~
tree −qmc −−l oad_pvalue \
−−a lpha [ alpha ] −−b eta 0 . 99 \
−i [ annotated t r e e ] −o [ output TOB]
~~~

The results of this tuning experiment are shown in Fig. 5. The estimated TOBs given as input to NetCS and NANUQ+ used the 3f1a option and significance threshold *α* = 10^−4^.

**Fig. 5:**
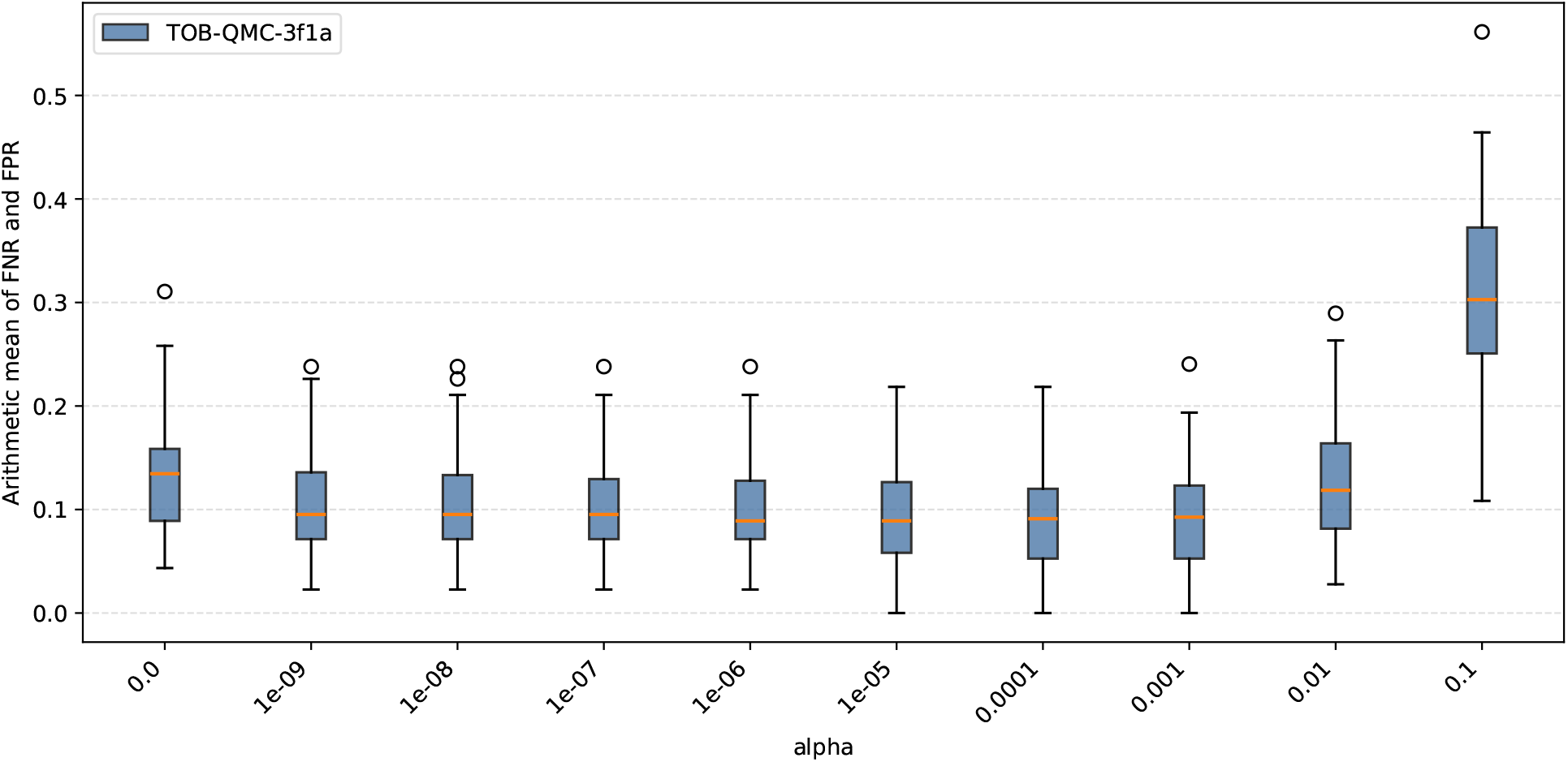
Experiment 3: Hypertuning parameters for TOB estimation. The *x*-axis is the significance threshold *α* required by TOB-QMC-3f1a. The *y*-axis is TOB topological error, specifically the arithmetic mean of FNR and FPR. FNR is the number of branches in the true TOB missing from estimated TOB divided by the number of branches in true TOB. FPR is the number of branches in the estimated TOB missing from the true TOB divided by the number of branches in the true TOB. Each boxplot has the 44 out of 50 replicates datasets for the 26-taxon model condition from the CAMUS study; these data sets are <u>not</u> used in experiment 3 beyond hypertuning. TOB-QMC failed to run on the 6 excluded replicates. Orange lines are medians.

**Fig. 6:**
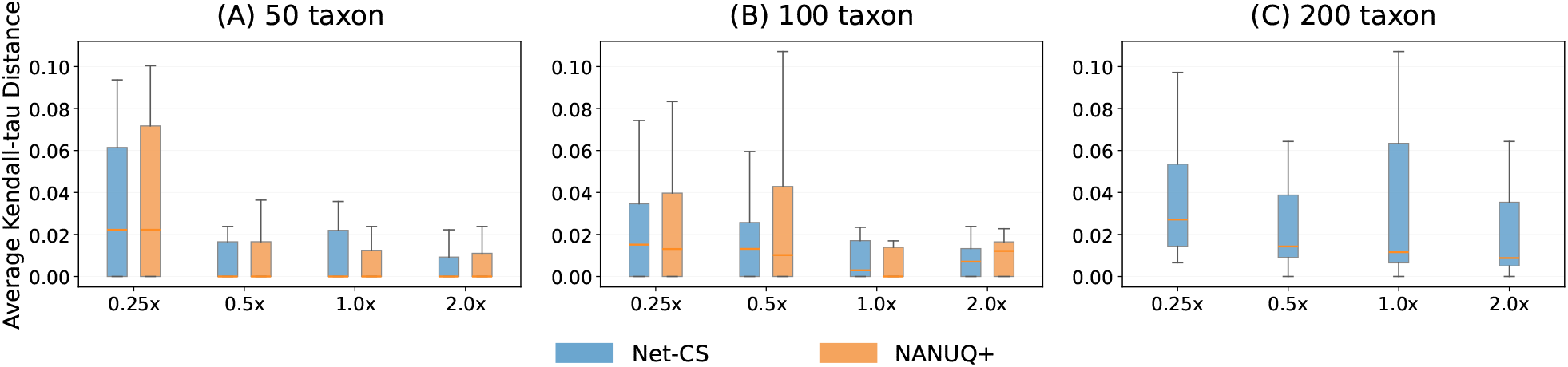
Experiment 2: Average Kendall-tau distance. Three subfigures (A), (B),(C) correspond to data sets with 50 taxa, 100 taxa, and 200 taxa, respectively. The *y*-axis is the average Kendall-tau distance, which evaluates the error in circular order reconstruction. The *x*-axis is the scaling factor: 0.25×, 0.5×, 1.0×, and 2.0× in order of decreasing ILS. Each colored box represents a method, either NetCS or NANUQ+. Both methods were given all estimated gene trees, up to 1000. For 200 taxon data, only Net-CS results are presented since NANUQ^+^ did not complete within the maximum wallclock time of 25 hours. Orange lines are medians. Outliers are not shown.

### B.5 CAMUS Command

CAMUS (v1.0.0-6-g3138bcf) was run with command:

~~~
camus −o [ output p r e f i x ] [ rooted base t r e e ] [ input gene t r e e s ]
~~~

where the rooted base tree was computed with ASTRAL-IV.

## C Supplemental Results

